# Tunability enhancement of gene regulatory motifs through competition for regulatory protein resources

**DOI:** 10.1101/2020.10.11.335018

**Authors:** Swetamber Das, Sandeep Choubey

## Abstract

Gene regulatory networks (GRN) orchestrate the spatio-temporal levels of gene expression, thereby regulating various cellular functions ranging from embryonic development to tissue homeostasis. Some patterns called “motifs” recurrently appear in the GRNs. Owing to the prevalence of these motifs they have been subjected to much investigation both in the context of understanding cellular decision making and engineering synthetic circuits. Mounting experimental evidence suggest that 1) the copy number of genes associated with these motifs vary, and 2) proteins produced from these genes bind to decoy binding sites on the genome as well as promoters driving the expression of other genes. Together, these two processes engender competition for protein resources within a cell. To unravel how competition for protein resources affect the dynamical properties of regulatory motifs, we propose a simple kinetic model that explicitly incorporates copy number variation (CNV) of genes and decoy binding of proteins. Using quasi steady-state approximations, we theoretically investigate the transient and steady-state properties of three of the commonly found motifs: autoregulation, toggle switch and repressilator. While protein resource competition alters the timescales to reach the steady-state for all these motifs, the dynamical properties of toggle switch and repressilator are affected in multiple ways. For toggle switch, the basins of attraction of the known attractors are dramatically altered if one set of proteins bind to decoys more frequently than the other, an effect which gets suppressed as copy number of toggle switch is enhanced. For repressilators, protein sharing leads to emergence of oscillation in regions of parameter space that were previously non-oscillatory. Intriguingly, both the amplitude and frequency of oscillation are altered in a non-linear manner through the interplay of CNV and decoy binding. Overall, competition for protein resources within a cell provides an additional layer of regulation of gene regulatory motifs.

## I. INTRODUCTION

Gene regulatory networks (GRN) consist of various molecular regulators such as Transcription factors (TFs) that interact with each other to perform a repertoire of functions in cells, ranging from embryonic development to tissue homeostasis [1–3]. Intriguingly, certain sub-networks in the GRN, are overrepresented. Such sub-networks are called motifs [4–6]. Due to the ubiquitous nature of gene regulatory motifs, a quantitative understanding of the dynamical behavior of them has concerned systems biologists and synthetic biologists alike [7–10]. While theorists have focused on mapping out the dynamic range of these circuits, experimentalists have sought to build them using a bottom-up approach. Such engineering approach adds two-fold value to the field; on the one hand, it furthers our understanding of the design principles of gene regulatory motifs inside cells, on the other, it helps in fabricating artificial circuits.

Most of these studies implicitly assume that gene regulatory motifs remain functionally isolated in a cell[11–13]. This assumption suffers from the complication that genetic motifs are always connected to many other genes inside a cell, which couples these motifs to various aspects of cell physiology. One such aspect is that protein products pertaining to genes of a motif bind promiscuously to a large number of decoy sites, as well as functional regulatory sites. Evidently, regulatory sites for a given gene may perform as decoy sites for another gene. For instance, well-known *E.coli* protein CRP has around 400 binding sites per genome copy [14]. The protein product of NF-kB, a gene associated with a eukaryotic oscillator, bind to a large number of sites across the genome; chip-seq data revealed approximately 20 thousand binding sites with around 500 of them being functional [15–18]. Moreover, copy number of genes within regulatory motifs can significantly vary in cells. For example, CNV of genome is widely prevalent in humans, as has been demonstrated in numerous studies [19–23]. Genes expressed on plasmids [24] or multiple identical copies on the chromosome [25–27], highly replicated viral DNA genes [28], etc. manifest key examples of CNV. Evolutionary analysis suggests that in some cases, whole-genome duplications led to lineage diversification in yeast [29]. The ubiquitous nature of CNV and decoy binding invokes competition for protein resources. A recent study explored the effect of gene copy number fluctuation [30] on the dynamical properties of well-known regultory motifs. On the other hand, several papers in the recent past have considered the effect of decoy binding [5, 12, 18, 31–35] in understanding the behavior of specific regulatory motifs. A quantitative understanding of how the interplay between CNV and decoy binding dictate the dynamical properties of gene regulatory motifs remains lacking.

To this end, we dissect the dynamical properties of three well-known gene regulatory motifs, i) Autoregulatory motif [36, 37], ii) Toggle switch [38–40], and iii) Repressilator [7, 10], in the presence of CNV and decoy binding. By utilizing the inherent separation of time scales of the system, we theoretically dissect this system [41]. Our investigation reveals that competition for finite protein resources alters the time to reach the steady-state for all of the three motifs. Moreover, toggle switch and repressilator exhibit qualitatively distinct dynamical behaviors; the basins of attraction of a toggle switch is dramatically altered when one of the two sets of proteins bind to decoys more frequently. This alteration though is suppressed as the copy number of toggle switch is ramped up. For repressilators, the interplay of decoy binding and copy number variation leads to either inhibition or abatement of oscillations depending on the parameter space. Moreover, the impact of CNV and decoy binding is rather complex on the amplitude of oscillations; in a context dependent manner the amplitude can either increase or decrease. We discuss the relevance of these results in the context of development and synthetic biology.

## II. RESULTS

To decipher the impact of CNV and decoy binding of protein products of genes pertaining to regulatory motifs on their dynamical properties, we consider three well-studied gene regulatory motifs, autoregulatory motif, toggle switch, and repressilator. In particular, we add *N*_*p*_ copies of the motif under investigation and *N*_*d*_ number of decoy binding sites in the system to study how the interplay of these two dictate the transient and steady-state behaviors of the motifs. We employ the so called pre-factor method, as developed earlier [41] and extend it in a congruous manner to achieve our goal. The analytical method is briefly illustrated in the next section; the details of the method are provided in the SI. Moreover, the parameter values chosen for the various models in our study lie in range with experimental measurements in Bacteria and viruses (list of all parameters used in numerical simulations can be found in the figure captions and in the supplementary material)). In the following section, we start with the autoregulatory motif owing to its simplicity.

### A. Auto regulation

Autoregulation is the simplest regulatory motif which also happens to be ubiquitous in cells [36, 37]. It comprises of a single gene, the product of which regulates its own expression, as shown in Fig. 1A. In this motif, the promoter transitions between two states which are defined by the promoter being bound/unbound to the protein; the rate of transition from the protein bound state to the empty promoter state is *k*_−_ while rate of switching from the empty promoter state to the protein bound state is *k*_+_. When the promoter is protein-bound, mRNA production commences at a rate *α*. The basal mRNA production rate from the empty promoter state is *β*. Each mRNA molecule is translated at a rate *σ* which leads to monomeric protein production. Protein monomers reversibly bind to each other to form a dimer with rate *κ*_+_ which can subsequently dissociate into monomers at rate *κ*_−_. mRNA and monomeric proteins degrade at rates *γ*_*m*_ and *γ*_*p*_ respectively. In our model, the dimeric proteins can bind to *N*_*d*_ number of decoy sites in addition to *N*_*p*_ number of promoters. We assume that the binding/unbinding rates of dimeric proteins to promoter and decoys are similar. Here it is assumed that dimeric proteins degrade at a rate much slower than all other reactions of the system, which is typically the case in *E.coli* (CITE). Here, the goal is to expound the effect of *N*_*p*_ and *N*_*d*_ on the various properties that characterize the dynamics of the autoregulatory motif. Previous studies have separately studied the impact of decoy binding [32] and CNV [30] of an autoregulatory motif on its input-output relationship. Hence, this model serves as a good starting point for analysis of more complex regulatory architectures. Using principle of mass action kinetics, the model can be described by a set of coupled ordinary differential equations (ODEs) characterizing the time evolution of average concentrations of relevant chemical species in the system (see Fig. 1B). The number of gene copies and decoys enter the description via the concentration of promoter and decoys; *N*_*p*_ and *N*_*d*_ are proportional to the total concentration of promoter sites, *d* (where *d*_*o*_ and *d*_*u*_ stand for concentrations of occupied and unoccupied promoter sites respectively), and decoy sites 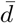 (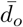 and 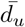 are concentrations of occupied and unoccupied decoy sites), where *N*_*p*_ = *d/C* and 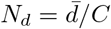; *C* = 10^9^*M* is the the molar concentration of a single molecule in the volume of an *E coli* cell, as reported earlier [30, 37]. To solve, these coupled ODEs, we make the simplifying assumption that the binding/unbinding of protein molecules to promoters and decoy sites equilibrates on a timescale which are comparatively faster than that of protein turnover. This assumption is consistent with numerous experimental [37] and theoretical studies [30, 41]. Following Bennett et al.[41], we use Quasi steady-state approximation which exploits this separation of time-scales in the system to solve for the transient and steady-state protein concentrations (see Fig. 1(C)). In particular, we keep track of the total monomer concentration. Details of the calculations can be found in the SI. This analytical procedure serves as a recipe for exploring the other motifs in the ensuing sections.

**FIG 1.**
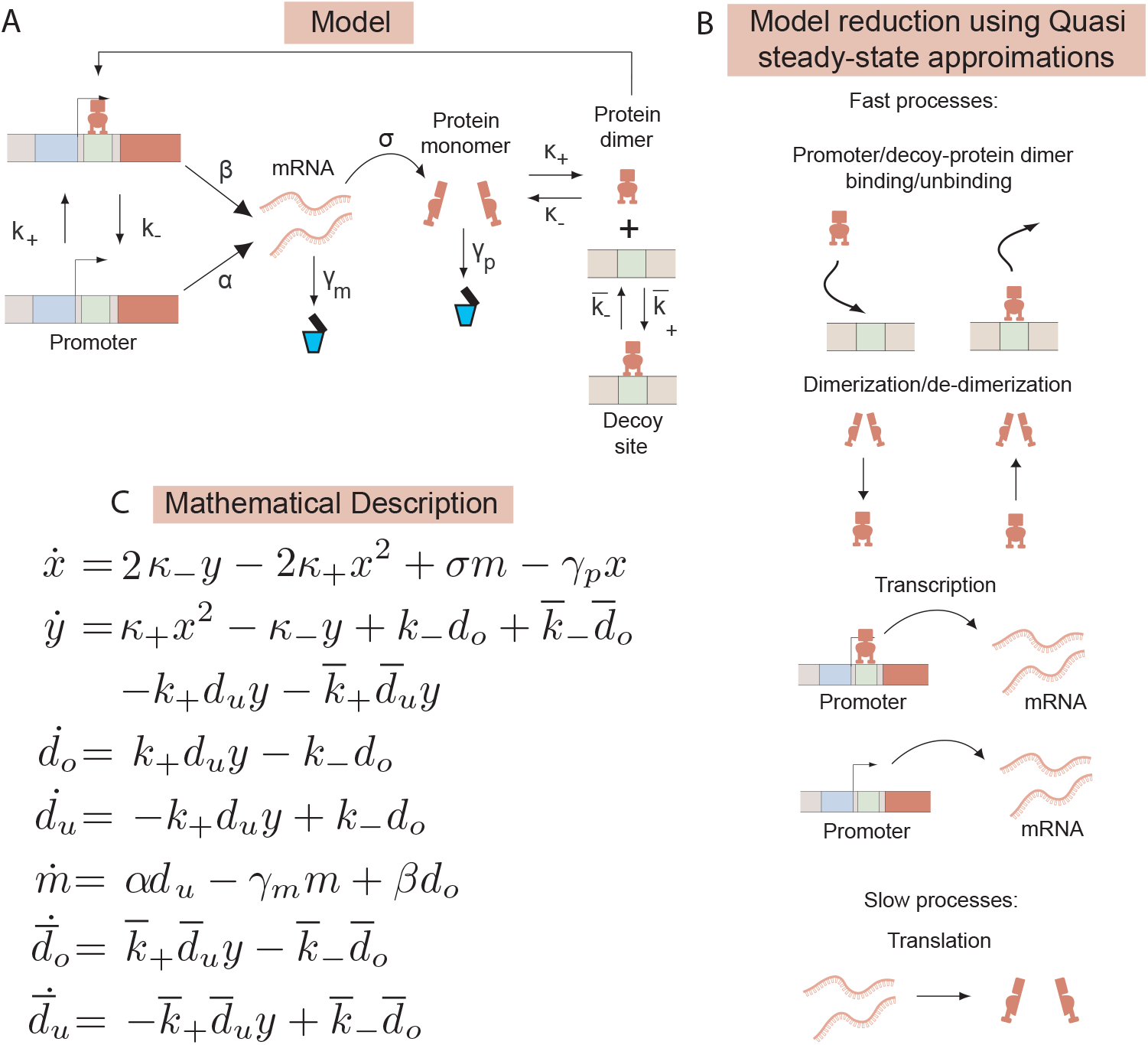
(A) Schematic representation of various processes for a positive autoregulatory motif in the presence of decoy sites. The promoter switches between a protein-bound and empty states with rates *k*_−_ and *k*_+_ respectively. Basal and regulated mRNA production rates are *α* and *β*. mRNA translation rate is *σ* which leads to monomeric protein production. Monomers reversibly bind to form a dimer with rate *κ*_+_ which subsequently can dissociate into monomers at rate *κ*_−_. mRNA and monomeric proteins degrade at rates *γ*_*m*_ and *γ*_*p*_ respectively. We assume that the dimeric protein can bind/unbind to promoter or decoys with the same rates. (B) Governing equations characterizing the time evolution of the relevant variables in the system. Variables refer to concentrations of protein monomers *x*, protein dimers *y*, unoccupied and occupied promoters *d*_*u*_ and *d*_*o*_, unoccupied and occupied decoys 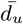 and 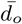, and mRNA *m*. (C) Separation of time scales, List of fast and slow processes we consider (mRNA and protein degradation rates are not shown).

Intriguingly, our model predicts that the mean expression level remains the same for varying number of decoy sites. However, time to reach the steady-state is enhanced as we increase the number of decoy sites (see Fig. 2A). As the number of decoy sites is enhanced, the possible number of configurations in the system increases, resulting in a higher amount of time to reach steady-state levels. This result is further confirmed by considering the time it takes to reach half the level of steady-state protein expression. This time is called the half-times. We find that in the absence of decoys, these half-times increase linearly with copy number *N*_*p*_ (see SI Fig. 1A). Moreover, half-times increase monotonically as a function decoy number and exhibit rapid increase for higher number of decoys. For smaller decoy numbers, half-times do not change in an appreciable manner, while for higher decoy numbers the change in half-times is significant. The point of rapid increase shifts to higher values of decoys as the gene copy number increases(see Fig. 2B). In other words, there exists a ‘tug of war’ between gene copies and decoys, which dictate the transient dynamics of autoregulation. It must be noted that the above-mentioned findings are a result of the prefactor method we employ. These results hold when we solve for the full system of equations, as shown in Fig. 2A,B) for a wide range of parameter values (see SI Fig. 3).

**FIG 2.**
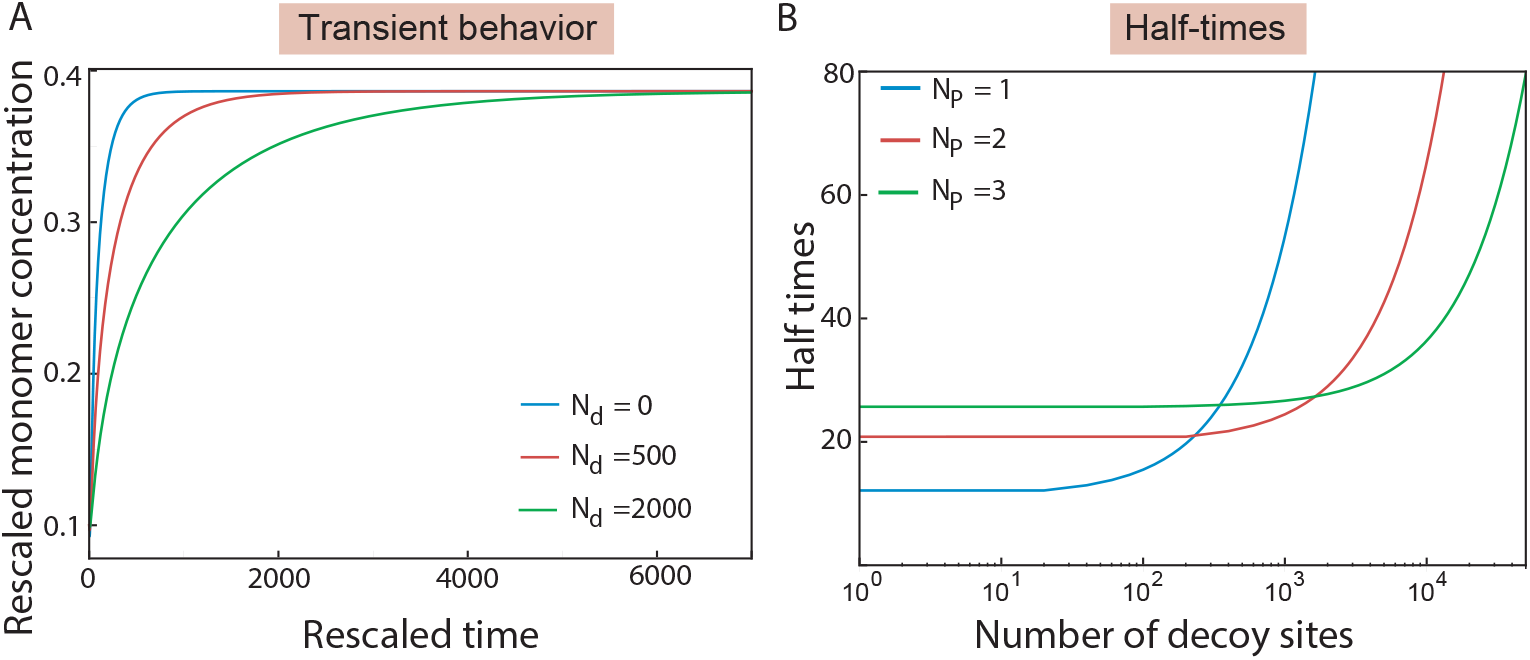
(A)Attainment of steady state of total monomer concentration. The presence of decoys affects the transient dynamics; the time it takes to reach the steady-state is enhanced as the number of decoys in increased (B) Half-times as functions of decoy copy numbers are shown for different copy numbers of the motif. Here half-times is defined as the time it takes to reach half of the steady-state monomer concentration.

**FIG 3.**
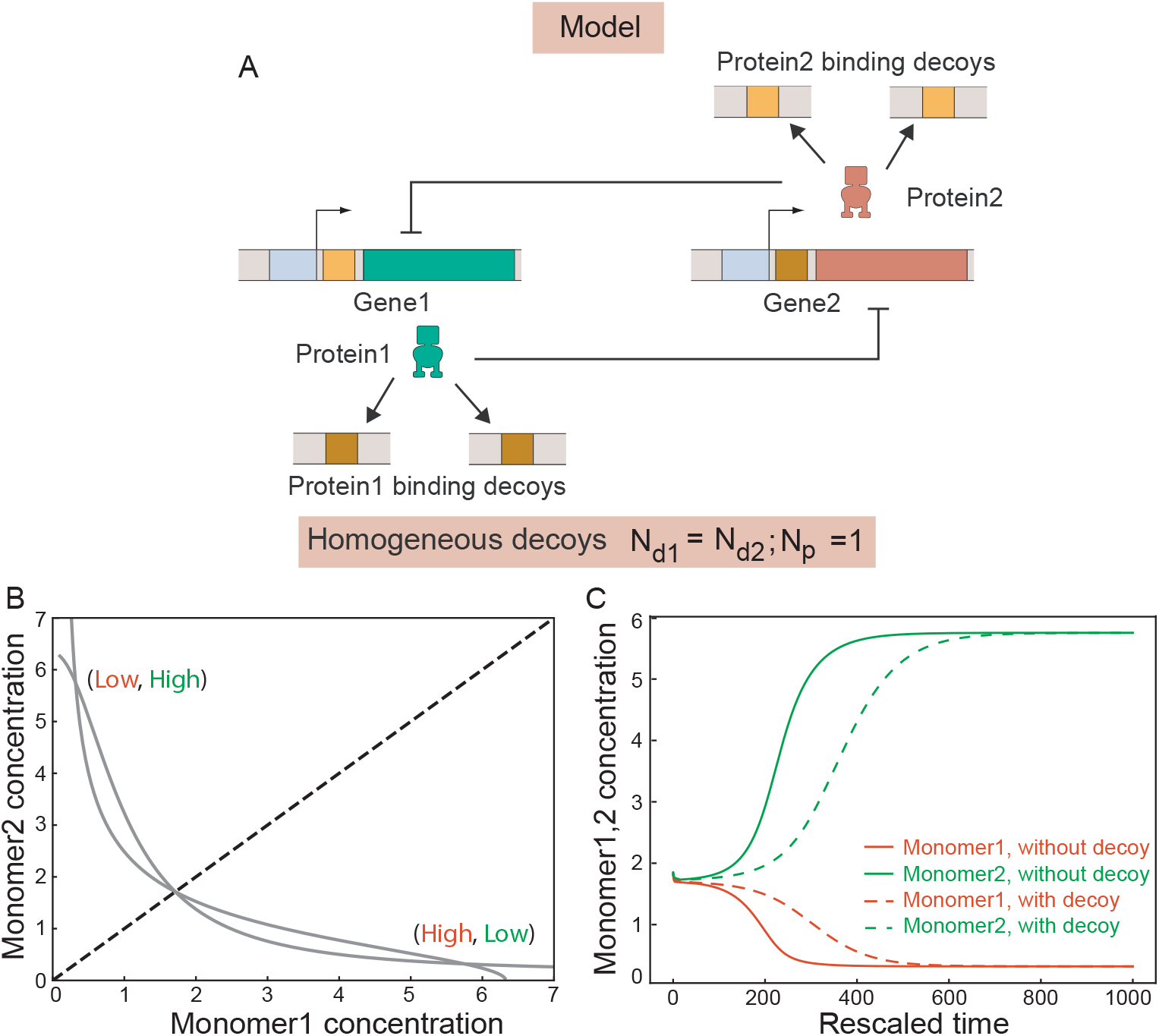
(A) Cartoon of the toggle switch. Two genes repress each other. The protein products pertaining to the genes can bind to decoys (B) Phase space of the toggle switch in presence of homogeneous decoys. The dashed line indicates a separatrix which defines the two stable states of toggle switch (High,Low) and (Low, High). The grey lines are nullclines for the toggle switch system which intersect at two stable and one unstable fixed points (see SI for details). (C) shows an example of the dynamics in the basin of attraction for (Low, High) state with and without decoys. Note that the presence of decoys increases the time to to reach the steady-state.

### B. Toggle switch

The toggle switch has attracted a lot of attention in recent past, since it is one of the simplest regulatory motifs that exhibit bistability [38–40, 42, 43]. This switch is often posited as the canonical model to characterize how multicellular organisms make “either/or-like” cell-fate decisions during development. Other instances of bistability include the lysis/lysogeny circuit of bacterio-phage lambda [44], the lac operon repressor system in *E. coli* [45, 46], cellular signal transduction pathways [47–50].

Toggle switch consists of two genes, that mutually repress each other, as shown in Fig. 3A. This system ex-hibits two stable equilibrium points (also known as attractors) in which one of the two genes is expressed at a high level while the other is expressed at a low level. In addition to these states, there is one unstable equilibrium point at which both the genes are expressed at some intermediate level. Associated with the two attractors are distinct regions of gene expression space of the two genes that are referred to as basins of attraction; a system evolved from an initial point within a basin of attraction eventually iterates into the corresponding attractor (see Fig. 3B). A straight line called the separatrix delimits the two basins of attraction. Our aim in this paper is to investigate if the presence of CNV and decoy binding affect these dynamical features, in particular the basins of attraction of toggle switch. In order to achieve this goal, we assume that in addition to *N*_*p*_ copies of the switch, there exist two sets of decoy sites (*N*_*d*1_, *N*_*d*2_ copies respectively) that the protein products of the two genes can bind to. The analysis is carried out by employing the same mathematical formalism as before (see SI for details).

First, we consider the scenario when copy number of switch is one and the two species of decoys are homogeneous i.e. number of both sets of decoy sites are equal with identical protein binding affinities. Under these conditions, the steady-state behavior of the toggle switch is not altered; the attractors and the corresponding basins of attraction remain identical to the usual toggle switch. However, the transient dynamics are impacted as in the autoregulatory motif in the presence of decoys. Time to reach the steady-state is slowed down substantially (see Fig. 3C) due to the system having to sample a larger space of possible microstates.

Next, we consider the scenario in which two species of decoys are heterogeneous; number of decoys pertaining to protein product of gene2 is higher than that of gene1. However, their protein binding affinities are equal. As shown in Fig. 4A, the basin of attraction of gene2 is substantially decreased at the expense of that of gene1. Let us consider the case in which in the absence of decoys, gene2 expression is high and expression of gene1 is low. Now, introducing a higher number of decoys corresponding to gene2 leads to its protein product on average getting occupied more often than that of gene1. This results in a reduction in the average level of repression gene2 exerts on gene1. Therefore, gene1 can switch to a higher level of expression and in turn, represses gene2. Thus, an initial point in the expression space of the two genes, that previously evolved to the first attractor now go to the second one, as shown in Fig. 4B. This effect remains intact when the two sets of decoys are identical in numbers but bind to the respective proteins with unequal affinities (see SI Fig. 1(B)).

**FIG 4.**
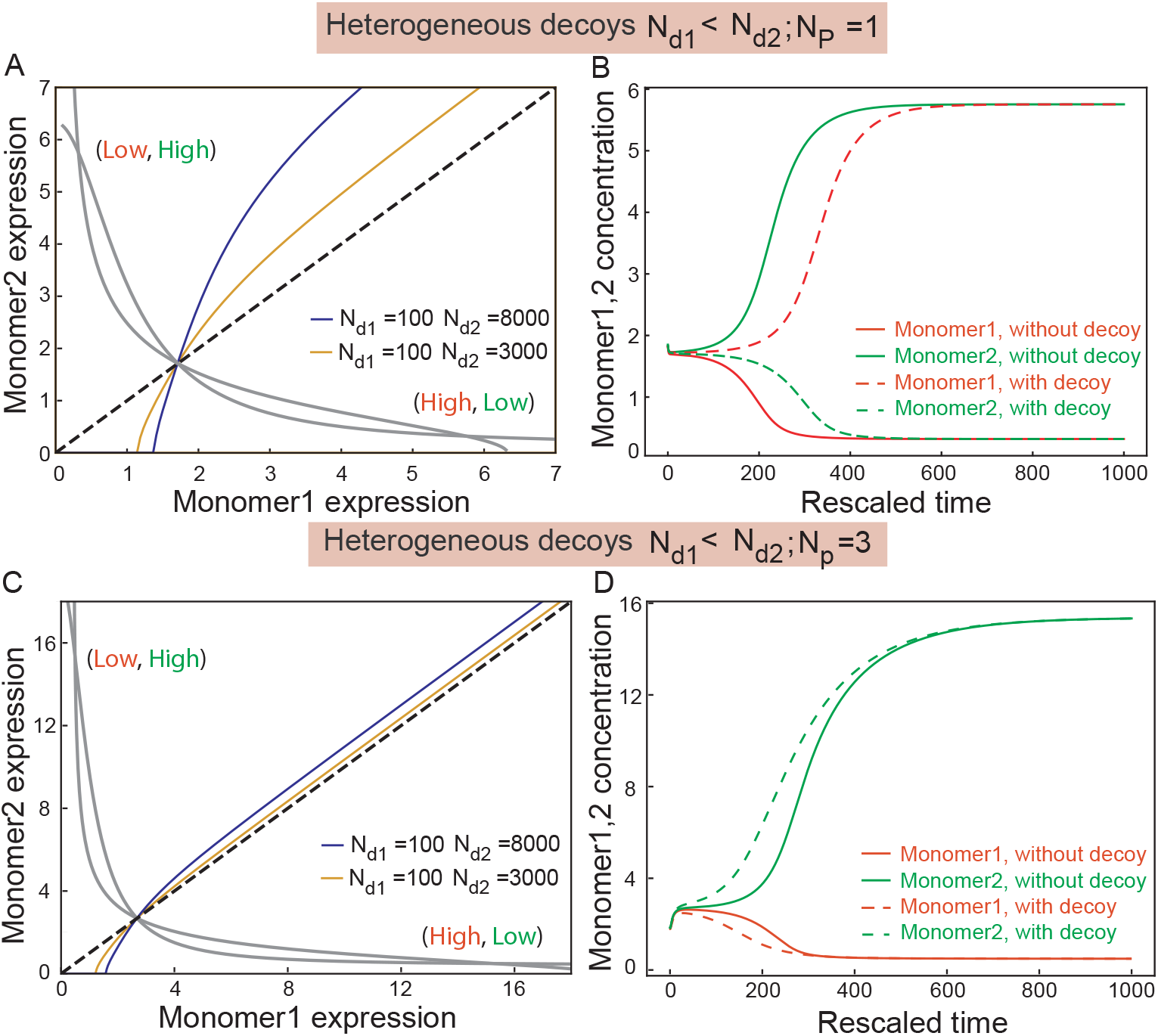
(A) Phase space of the toggle switch in the presence of heterogeneous decoys i.e. when the numbers of decoys corresponding to two genes are different (number of decoys for protein2 is higher than the number for protein1). The separatrix begins to exhibit a curvature that changes with the degree of dissimilarity in decoy numbers. The basin of attraction associated with (high,low) state thus expands at the expense of the one related to (low,high) state. The grey lines are nullclines for the toggle switch system which intersect at two stable and one unstable fixed points (see the supplementary material for details). (B) The example of Fig. 2B is re-plotted in the presence of dissimilar decoys. Initial conditions that evolved to (low,high) state earlier evolved to (high,low) state. (C) Phase space of the toggle switch for *N*_*p*_ = 3 copies of the switch in presence of heterogeneous decoy binding. Increase in gene copy number rescues the effect of heterogeneous decoy binding; the effect of decoys is suppressed. It must be noted that the average expression level of both the genes are higher for when the copy number of switch is higher. (D) Such rescue effect is further evident from the transient behavior of the system; time to reach the steady-state is reduced.

**TABLE 1.** Summary of observations for the three motifs with change in promoter and decoy copy numbers.

| Motif | Observations |
| --- | --- |
| Autoregulator | Change in promoter concentration influences the steady state levels of the gene.<br>Increase in decoy concentration increases half-times to attain the steady state level. |
| Toggle Switch | Asymmetry in binding affinities to decoy sites alters the basins of attraction corresponding to the stable states. Increase in promoter concentration counterbalances the effects introduced by the increase in decoy sites. |
| repressilator | Increase in promoter concentration expands the oscillatory regime in the stability diagram.<br>Change in decoy concentration leads to emergence of a new oscillatory regime while frequency and amplitude are affected in a nonlinear fashion. |

A key result of our paper is that the effect of heterogeneous decoy binding is significantly suppressed when the copy number of toggle switch is enhanced. As shown in Fig. 4C, the basins of attraction can be rescued at high copy number. Consequently, points in the basin of attraction of one attractor that switched to the other due to heterogeneous decoys returns to the former attractor when the copy number is increased (see Fig. 4D).

As such, our theory makes specific predictions about how an interplay between CNV and decoy binding alters the dynamical properties of toggle switch.

### C. Repressilator

The repressilator typically consists of three genes coupled in a cyclic structure, such that each gene represses the next (see Fig. 5A), generating oscillations in protein concentrations [7, 10]. There exist numerous examples of naturally-occurring repressilators [51]. A classic example is that of the circadian clock which oscillates according to the day-night cycle [52, 53]. In mammals a set of three genes, cryptochrome (Cry), period (Per), and Rev-erb serve as a major core element of the circadian network [54]. A repressilator is responsible for controlling circadian timing in A. thaliana Synthetic circuits built from well-characterized genetic parts can also exhibit oscillations [55, 56].

**FIG 5.**
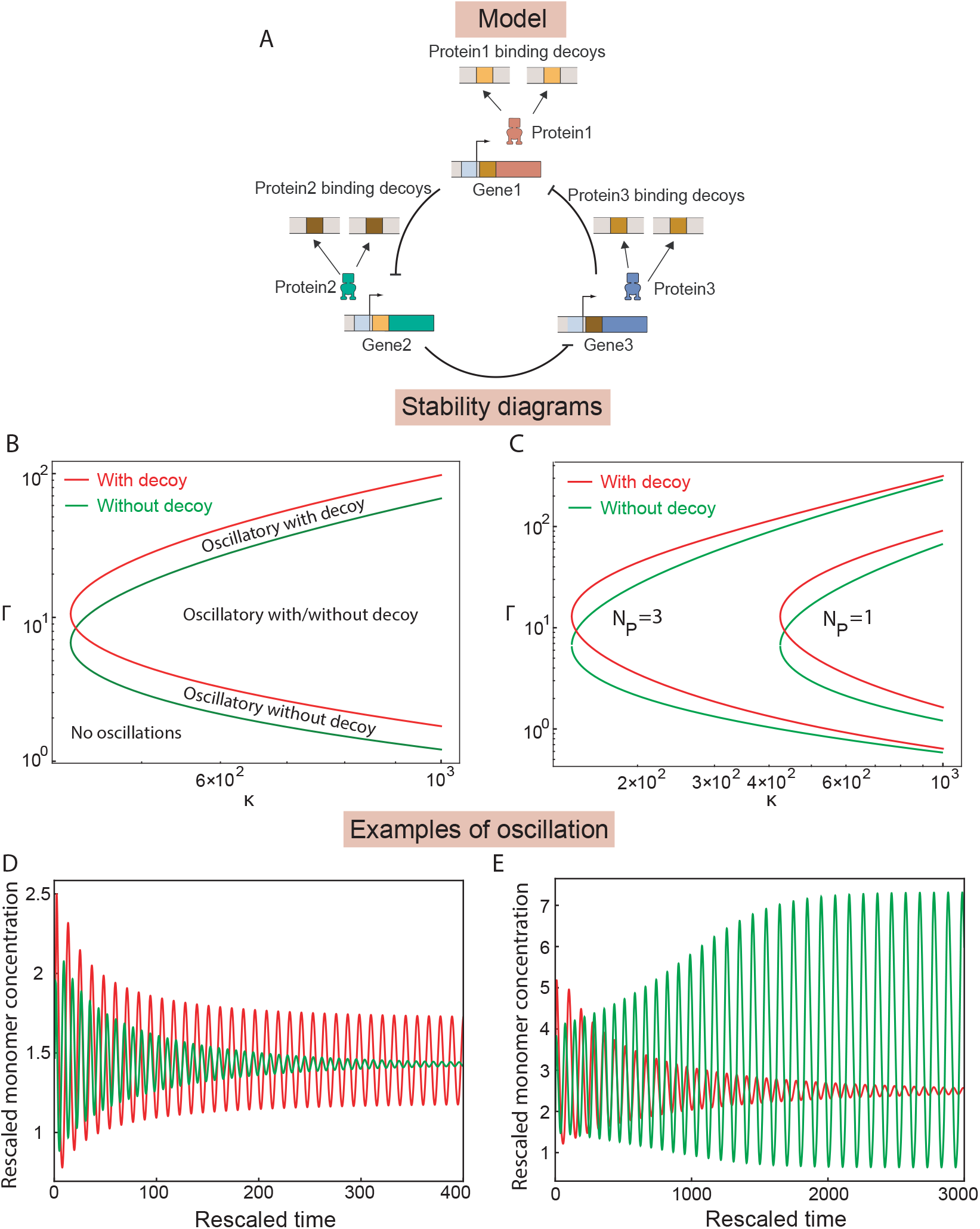
(A) Cartoon of the repressilator. Three genes repress each other in a cyclic structure. The proteins produced from these genes bind to decoy sites. (B) Stability diagram in Γ − *κ* parameter space, where Γ = *γ*_*p*_*/γ*_*m*_ and *κ* = *ασ/*(*γ*_*m*_*γ*_*p*_) (see Fig. 1C and supplementary material for details of the parameters). The blue curve is for the case without decoys while the red curve indicates the case when decoys are added. (C) We show stability diagrams for *N*_*p*_ = 1, 3 with decoy number *N*_*d*_ = 1000. A part of the steady state regime becomes oscillatory in the presence of decoys and vice versa. (D) An example of dynamics from the region exhibiting oscillatory behavior without decoys becomes steady in the presence of decoys. (E) A complementary example from the region for the opposite case is shown.

As a dynamical system, repressilator exhibits stable limit cycles appearing due to a supercritical Hopf bifurcation of the stable equilibrium point [30, 41]. Consequently, the system exhibits oscillations for a wide range of parameters. In other words, the protein products of each of the three genes in the circuit oscillate in time. The corresponding bifurcation diagram displays a separatrix demarcating the oscillatory from the steady-state regime. We are interested in unraveling the consequences of CNV and decoy binding of proteins expressed from these genes upon the dynamical behavior of repressilator.

To carry out such an analysis, we consider the number and protein binding affinity of the three sets of decoy sites corresponding to three kinds of proteins products in the system to be identical. Numerical simulations show that the copy number of motif and decoys significantly alter the oscillatory regime. In particular, our analysis evinces that a considerable region in the parameter space that exhibited steady-state behavior turns oscillatory in presence of CNV and decoy sites, as evident from the standard bifurcation diagram (see in Fig. 5B). In addition, there emerges a parameter regime which is no longer oscillatory (for example, see Fig. 5C and Fig. 5D).

We further study the impact of CNV and decoy binding on the frequency (also known as Hopf frequency) and amplitude of oscillation. An expression of frequency *ω* has been derived analytically which involves gene copy number *N*_*p*_ and decoy number *N*_*d*_ (see the SI for details). We seek to explore how this frequency changes as a function of *N*_*p*_ and *N*_*d*_. In particular, frequency ratio *ω*_*r*_ is defined as the ratio of *ω* and *ω*_0_, which is the frequency for *N*_*p*_ = 1 and *N*_*d*_ = 0. Frequency ratio *ω*_*r*_ monotonically decreases with increasing decoy number. While the overall frequency is further reduced as the gene copy number is increased, the rate of decrease in frequency ratio is smaller for higher gene copy number, as shown in Fig. 6A. Once again, the influence of decoys is seen to be negated by CNV.

**FIG 6.**
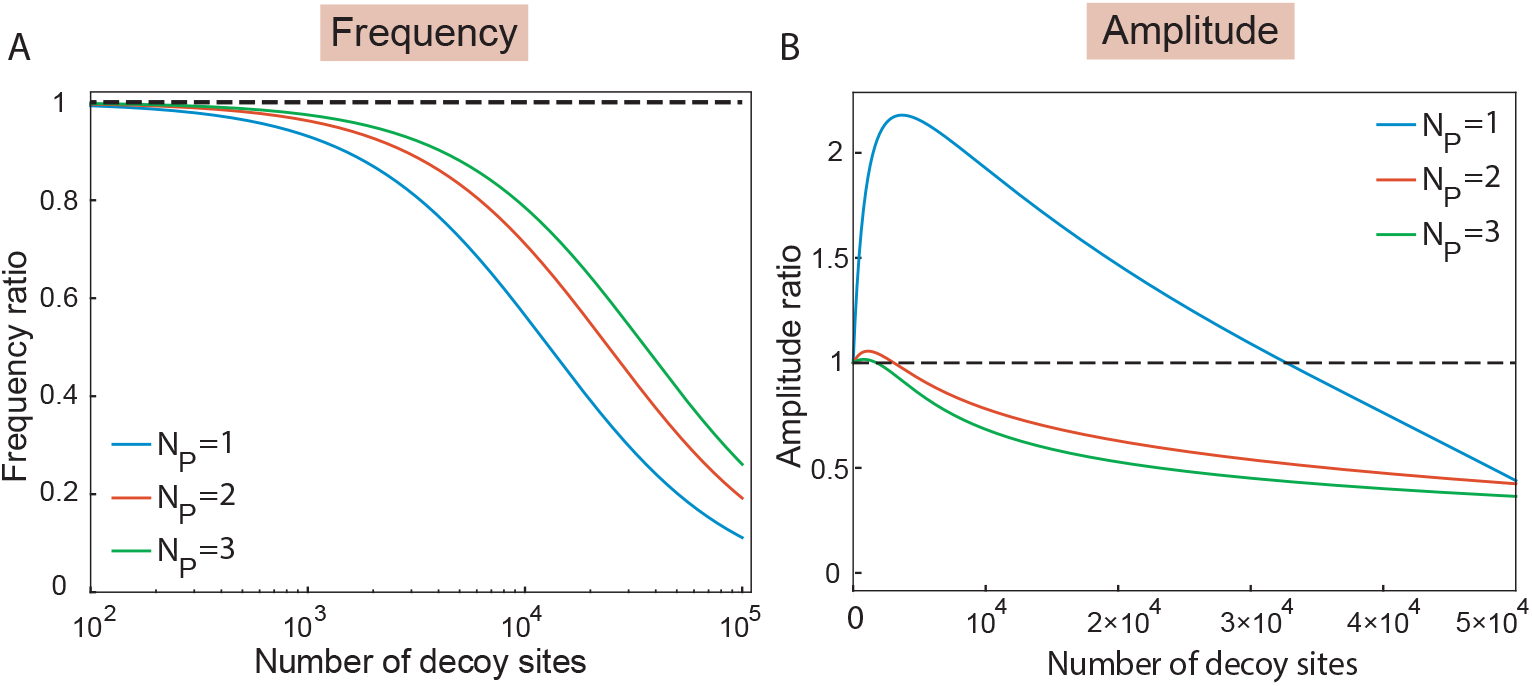
(A) Frequency ratio of oscillation with and without decoy binding is shown. Ratios for three different gene copy numbers are shown. The dashed line in black indicates *ω*_*r*_ = 1 for reference. (B) Amplitude ratios with and without decoys for different copy number of genes are shown. It shows a nonlinear behavior as a function of decoy number; initially it does not vary much but then shows a hump followed by rapid decline as decoy number is increased. The dashed line in black indicates *A*_*r*_ = 1 for reference.

Interestingly, we discover that the amplitude of oscillation demonstrates both an increase and decrease in presence of decoy binding sites. We define an amplitude ratio *A*_*r*_ (similar to *ω*_*r*_). The ratio is stable for small *N*_*p*_ and *N*_*d*_. A hump in the curve appears for moderate *N*_*d*_ for *N*_*p*_ = 1 which indicates the region of high amplitude followed by a rapid decline for high *N*_*d*_. With higher gene copy numbers, the hump begins to disappear and subsequent fall in the ratio is relatively sharper as shown in Fig. 6B. This behavior can be further confirmed by observing the monomer concentration as a function of the number of decoy sites, as shown in (see SI Fig. 4). It must be noted that for a large *N*_*d*_, the repressilator moves from the oscillatory to the steady-state regime. This behavior is sensitive to various parameters of the motif such as monomer degradation rate, protein binding affinity of decoys etc. (see SI Fig. 1(C)).

## III. DISCUSSION

In this manuscript, we have investigated how gene copy number variation and presence of decoy sites together modulate the dynamical properties of three well-known regulatory motifs. Copy number variation of genes and addition of decoy sites invoke a competition between the promoters and decoys for the pool of regulatory proteins in the system [30, 33, 34]. Such resource sharing couples the dynamics of genetic circuits with cell physiology in general, and provides a rich context in which these circuits operate in cells [57]. This is particularly pertinent in engineering synthetic circuits, which entails a firm quantitative understanding of how the various contexts in a cell impacts the circuit behavior [35, 58]. Recent studies in the field of regulatory biology have started exploring the impact of resource sharing on the dynamics of genetic circuits [30] and single genes [5, 12, 18, 31–35, 59, 60]. However, a comprehensive theoretical understanding is still lacking. In this manuscript, we use the pre-factor method, as introduced earlier [41], to study the transient and steady-state properties of the motifs under investigation.

Our study reveals that for toggle switch, the basins of attraction of the two attractors can be significantly altered in face of decoy binding. Similar results were obtained previously, when Lyons et al. [12] showed that the addition of a downstream component or load to the toggle switch alters the underlying potential energy landscape. The downstream component in that case could be a protein or a small molecule such that the bound complex prevents one kind of repressor from binding to and repressing its conjugate promoter. In effect, our system invoked the same effect of altering the effective concentration of one of the proteins by occupying them with more decoy sites. Intriguingly, we find that a relatively smaller increase in gene copy number can dramatically suppress this effect. This result implies that using multiple copies of synthetic circuits in a cell can keep the dynamics of these circuits relatively insulated. Moreover, the findings in this paper may have important biological implications. For instance, toggle switch is often invoked as a minimal model of cellular decision-making during mammalian development, where cells need to choose between alternate fates. Often cell-types associated with these fates need to be produced in different proportions. Our results suggest an exciting possibility of transcription factor resource sharing being a potential mechanism for controlling the proportion of cells belonging to different cell-types during development.

Introduction of decoy sites impact the oscillatory behavior of a repressilator in multiple ways. The stability diagram gets altered significantly as CNV and decoy binding are employed. As such, both these features increase the parameter space of oscillations. Amplitude and frequency of oscillations can be widely tuned as a function of the number of decoy sites and gene copy number. The impact of decoy binding are consistent with a previous study that explored the impact of decoy binding on NF-kB network which exhibits oscillation [18]. Although the concomitant circuit architecture was different, they observed an alteration in amplitude and period of oscillations. On the one hand, our results provide constraints on how to build a synthetic repressilator, on the other, the results imply enhanced tunability of the circuit’s dynamical behavior. Whether such differential control is functionally utilized by cells is an intriguing question.

It must be emphasized that the pre-factor method we employ to solve for the transient and steady-state properties of the system suffers from notable limitations. This method is a quasi-steady-approximation with an appropriate correction factor that appears due to the correct identification of the truly slow variable in the system. Such a reduction scheme heavily relies upon the fact that timescale separation exists in the system. If different chemical species of the system react with similar timescales i.e. are not separable into sets of slow and fast reactions then this reduction method is not applicable and our results would not hold. Moreover, the method ignores the stochastic fluctuations introduced due to the intrinsic randomness of biochemical reactions present in the system [41].

Overall, our study evinces that CNV and decoy binding are in a tug of war in terms of governing the behavioral properties of genetic motifs. Changes in gene copy number and decoy number can potentially function as tuning knobs within a nonlinear dynamical system as another layer of regulation.

## IV. ACKNOWLEDGMENT

All the codes for the computational analysis and generating the subsequent plots can be found on Github. Moreover, the codes were written in python. SD is grateful to Prof. Dr. Arnd Bäcker for his generous support during this work.

## Supplementary Information

### I. AUTO-REGULATION

We begin with the auto-regulatory motif; the protein product of a single gene regulates its own expression. Consider a simplified setting wherein protein monomers dimerize to create dimers which can bind/unbind with available promoters and decoys. Promoter bound protein dimers leads to transcription while decoy bound dimers remains inert. A detailed description of the model is given in the main text in Fig. 1A.

The dynamics of the system is captured by the following set of coupled differential equations:

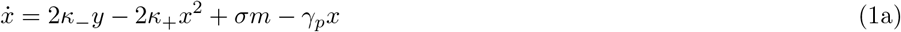

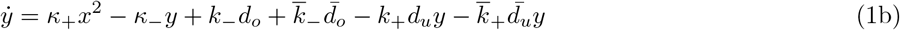

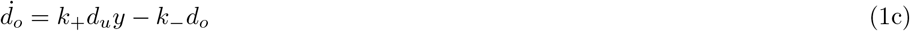

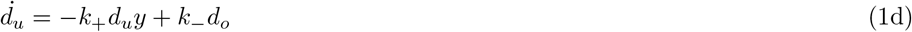

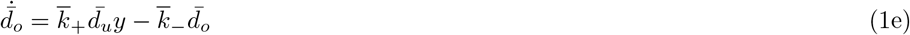

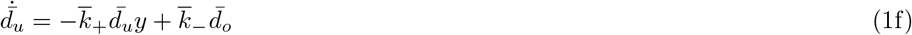

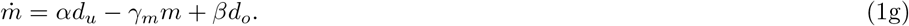

There equations represent the time evolution of the average concentration of following seven species -

*x* ≡ protein monomers

*y* ≡ protein dimers

*m* ≡ mRNA

*d*_*u*_ ≡ unoccupied promoter sites

*d*_*o*_ ≡ occupied promotor sites

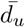 ≡ unoccupied decoy sites

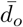 ≡ occupied decoy sites

The parameters involved are as follows:

*σ* ≡ translation rate,

*α* ≡ basal transcription rate,

*β* ≡ regulated transcription rate

*γ*_*m*_ ≡ degradation rate of mRNA,

*γ*_*p*_ ≡ degradation rate of protein monomers

*k±* ≡ binding and dissociation rates of the protein to a promoter,

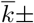 ≡ binding and dissociation rates of the protein to a decoy site,

*κ±* ≡ binding and dissociation rates of the protein to themselves.

We have the constraints that total amount of promoters and decoys remain conserved:

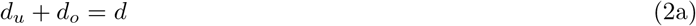

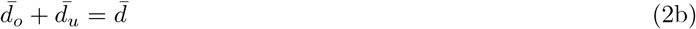

where, *d* and 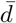 are total concentrations of promoter sites and of decoy sites respectively. We substitute these expressions into Equation (1) and use the recalling that *d* = *N*_*p*_*C* and 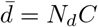. Here *N*_*p*_ and *N*_*d*_ are promoter and decoy copy numbers respectively. *C* = 10^−9^ Molar is the conversion factor denoting the concentration of a single molecule in the volume of an *E. coli* cell [1, 3]. Moreover, the degradation of dimers is ignored here for analytical tractability.

#### A. Multiple time-scale analysis

The governing Eqs. 1 are a set of seven dimensional nonlinear coupled differential equations. But, as described in several previous studies [1, 3], the system exhibits multiple time-scales which may be used to reduce the effective dimensions of the system which leads to simplification subsequent mathematical and computational analysis.

We assume that the dimerization and regulatory binding process are fast compared to transcription, translation, and degradation. These assumptions hold for a large number of bacterial and viral genes [1, 3]. Therefore, Eqs. 1(b)-1(f) involves fast processes while Eqs. 1(a) and 1(g) are slow. Consequently, the fast reactions will reach their steady-states earlier compared to their slow counterpart. By setting derivatives of *y, d*_*u*_, *d*_*o*_, 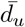, and 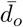 to zero, we obtain:

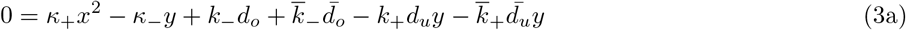

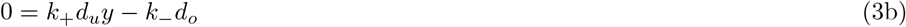

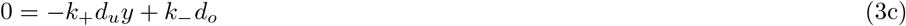

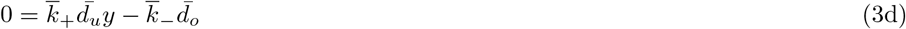

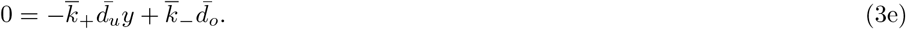

Thus, we obtain the steady-state values as below:

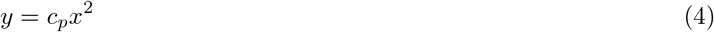

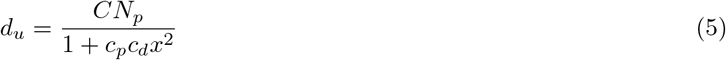

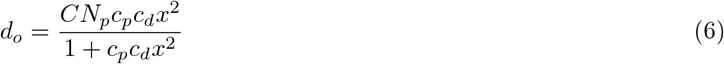

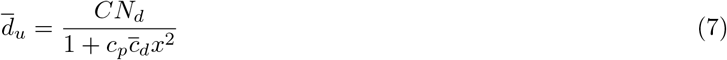

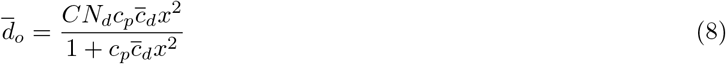

where, 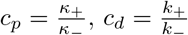, and 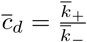.

These steady-state values can now be substituted in Eqs. 1(a) and Eqs. 1(g), to obtain:

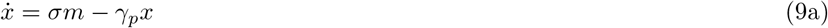

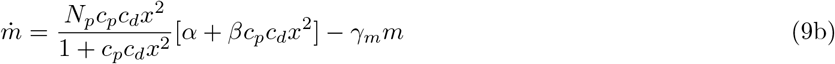

The continuous approximation scheme described above is called the quasi-steady state approximation (QSSA) [1–3] and is popular among theorists to model the dynamics of network motif. However, in order to track the transients correctly, an important correction in the form of a pre-factor was proposed [2]. We find the pre-factor method naturally allows for inclusion of copy number variation and presence of decoys as described below.

#### B. The pre-factor method

We note that reaction 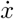 is actually a mixture of slow (translation and degradation) and fast reactions (dimerization and disassociation). Therefore, a *true* slow variable is the total concentration of protein molecules (in any form):

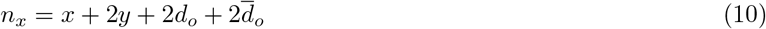

Time evolution of *n*_*x*_ is given by:

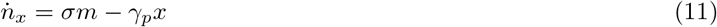

Now, substituting steady values in Eq. 10:

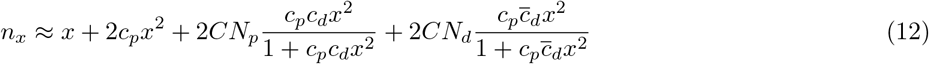

We can now express 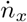 as follows:

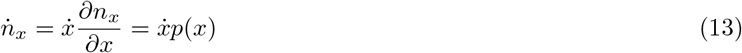

where, pre-factor *p*(*x*) is given by:

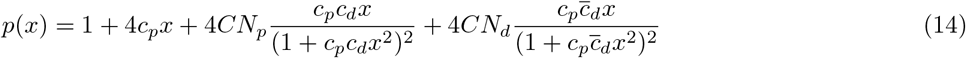

A new system of equations for time evolution of *x* and *m* is obtained using Eq. 11 and Eq. 14:

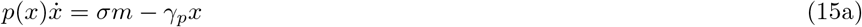

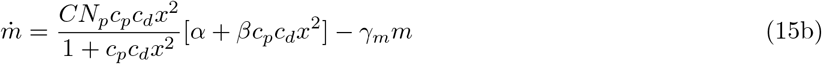

Note that any term involving decoy sites does not appear explicitly in the Eq. 15. Their influence is buried in the pre-factor *p*(*x*) (see Eq. 14).

#### C. Comparison of the pre-factor method with the full system

On comparing the results obtained form pre-factor method and those from the full system (without any reduction scheme used), we find that

1. Both the methods lead to qualitatively similar dynamics.
2. Increasing decoy sites results in longer transients in the full system. This behavior once again is captured by the pre-factor method.

The corresponding plots are given in SM Fig. 3 at the end.

#### D. Computation of Half-life *T*_1*/*2_

We define *T*_1*/*2_ as the time the system takes from an initial concentration of proteins (and mRNA) to reach one-half of its steady state value. Figure 2 in the main text shows the effect of decoys on *T*_1*/*2_ when *N*_*p*_ = 1.

To determine a reliable value of *T*_1*/*2_, we consider an ensemble of 100 random initial conditions drawn from a uniform distribution in a neighborhood Δ ≈ 10^−2^ of the stable equilibrium point. For a given *N*_*p*_, *T*_1*/*2_ averaged over the ensemble is then taken to be the value of half-life. In the absence of decoys, the computed values for various *N*_*p*_ ∈ {1, 2, 3, 4, 5, 6} is plotted in Fig. 1 (A). A similar procedure is used to compute *T*_1*/*2_ in the presence of decoy sites shown in Fig. 2 of the main text.

#### E. Parameters

We use the following parameters values to perform computations [1–3]:

*C* = 10^−9^*M, c*_*p*_ = 10^7^*M* ^−1^, *c*_*d*_ = 10^7^*M* ^−1^, 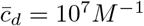, *σ* = 0.5.

|  |  |
| --- | --- |
| Time evolution of monomer concentration and half time computations of Figs. 2(A-B) were done using the following model parameters: | $\alpha = 0.025 \text{ min}^{-1}, \beta = 1.3 \text{ min}^{-1}, \gamma_p = 0.05 \text{ min}^{-1}, \gamma_m = 0.10 \text{ min}^{-1}$ [1–3]. |

### II. TOGGLE SWITCH

A toggle switch motif contains 2 genes with two possible states of expression – (Low, High) and (High, Low) which are obtained by one gene partially inhibiting transcription of the other. We assume regulated transcription rates *β*_1,2_ to be smaller than basal transcription rates *α*_1,2_ i.e. *β*_1,2_ *< α*_1,2_. For simplicity, we keep *β*_1_ = *β*_2_(= *β*), *α*_1_ = *α*_2_(= *α*), and *γ*_*p*,1_ = *γ*_*p*,2_(= *γ*_*p*_).

We use Eqs. 15 and the pre-factor (see Eq. 14) to write the model for the dynamics of the toggle switch, as follows:

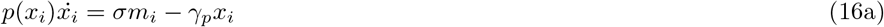

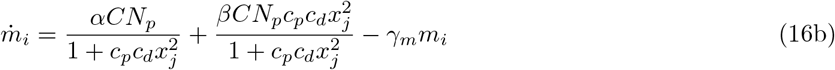

where *i* ∈ {1, 2}, *j* ∈ {2, 1}.

We use following rescalings to simplify Eqs. 16, 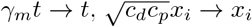, and 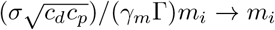, to obtain:

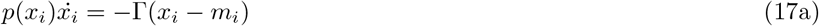

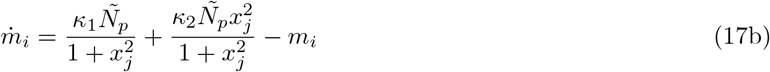

where Γ = *γ*_*p*_*/γ*_*m*_, *κ*_1_ = *ασ/*(*γ*_*m*_*γ*_*p*_), *κ*_2_ = *βσ/*(*γ*_*m*_*γ*_*p*_), 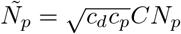, and *i* ∈ {1, 2}, *j* ∈ {2, 1}.

The pre-factor becomes:

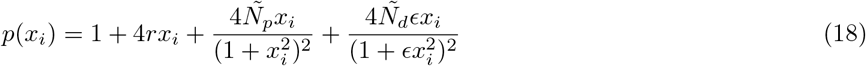

where, 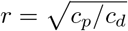 and 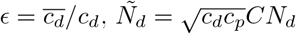. Also, note that the assumption *β < α* translates to *κ*_2_ *< κ*_1_.

#### A. Breaking of symmetry of decoys

Eqs. 17 consists of four equations as follows:

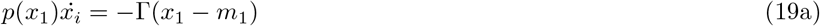

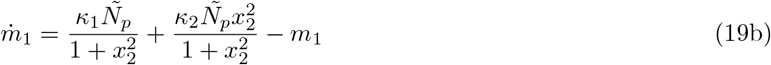

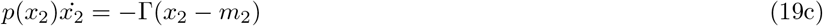

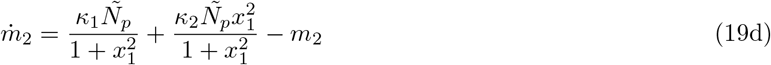

The pre-factors are:

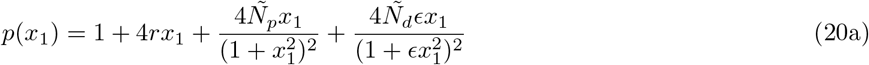

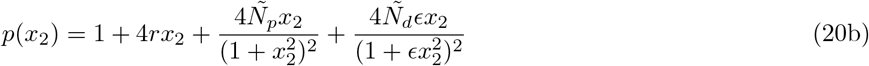

The equations above, we assumed that rate of binding and unbinding of protein dimers to decoys sites to be equal for both the genes, 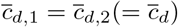, which implies *ϵ*_1_ = *ϵ*_2_(=*ϵ*).

Now, this symmetry of decoy sites is effectively *broken* by including two different species of decoys such that *N*_*d*,1_ *≠ N*_*d*,2_. It modifies pre-factors in Eqs. 20 as follows:

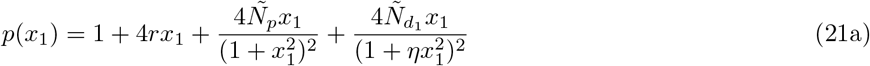

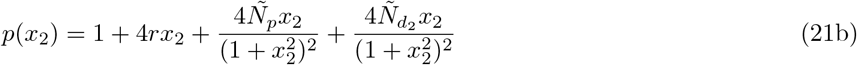

For simplicity of analysis, we assume *ϵ*_1_ = *ϵ*_2_ = 1. The main text shows the influence of *Ñ*_*d*,1_ *< Ñ*_*d*,2_ on the basin of attraction of the two stable fixed point at saddle node bifurcation.

##### 1. Basin of attraction for different protien binding affinities of decoys

As noted in the main text, an alternative way to break the symmetry of decoys is to consider identical decoy number with different protein binding affinities *ϵ*_1_ *≠ ϵ*_2_. We find the separatix sensitive to this change and as a consequence, basin of attraction of stable states (High, Low) and (Low, High) demonstrate considerable variation (see Fig. 1 (B)). This appears to be a non-trivial behavior and warrants deeper investigation; we leave it for future research.

#### B. Parameters

We use the following parameter values to perform computations:

*C* = 10^−9^*M, c*_*p*_ = 10^7^*M* ^−1^, *c*_*d*_ = 10^7^*M* ^−1^, 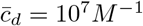, *σ* = 0.5.

|  |  |
| --- | --- |
| Basin of attraction and time evolution of monomer concentration as in Fig. 3,4 | $\alpha \approx 6.32 \text{ min}^{-1}, \beta \approx 0.14 \text{ min}^{-1}, \gamma_p = 0.05 \text{ min}^{-1}, \gamma_m = 0.10 \text{ min}^{-1} (\Gamma = 0.5, \kappa_1 = 632.50, \text{ and } \kappa_2 = 13.75). [1-3]$ |

### III. REPRESSILATOR

We use Eqs. 15 with *β* = 0 (no transcription from bound promoter sites) to get the model equations for the repressilator:

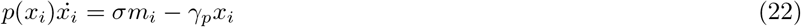

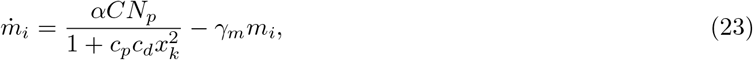

where *i* ∈ {1, 2, 3}, *k* ∈ {3, 1, 2}.

Rescalings: *γ*_*m*_*t* → *t* 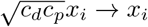, and 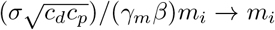, to obtain:

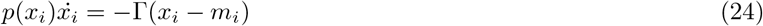

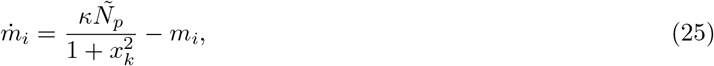

where Γ = *γ*_*p*_*/γ*_*m*_, *κ* = *ασ/*(*γ*_*m*_*γ*_*p*_), 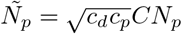.

The pre-factor is given by:

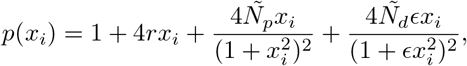

where 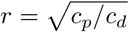 and 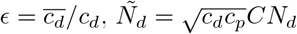.

#### A. Stability analysis and Hopf frequency

The repressilator defined by Eqs. 24 has a symmetric equilibrium 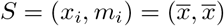 which is unique real solution of the equation:

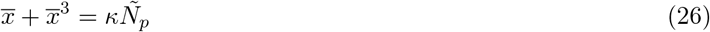

Jacobian of the system around this equilibrium reads:

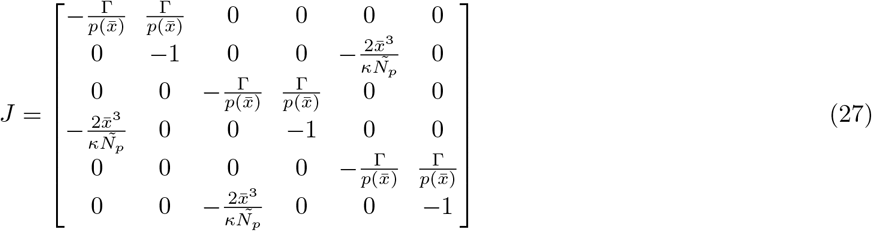

The Jacobian *J* has the following eigenvalues at the symmetric equilibrium defined above:

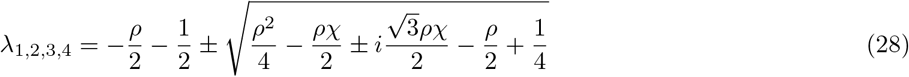

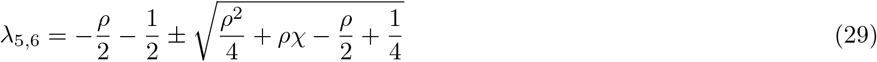

where 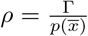 and 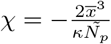.

##### 1. Bifurcation analysis: Hopf Frequency

When a complex eigenvalue *λ* cross the imaginary axis, we get the Hopf Bifurcation. At the point of bifurcation, the real part of an eigenvalue becomes zero and the imaginary part gives the frequency of oscillations.

Taking 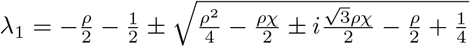

We write *λ*_1_ as follows:

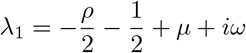

Setting the real part to zero, we get:

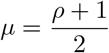

But, *µ* + *iω* is given by:

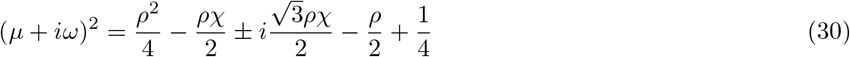

Equating the real and imaginary parts, we find, for the imaginary part:

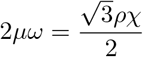

Substituting the expression for *µ*,

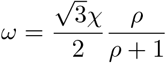

Substituting the expressions for *ρ* and *χ*, this imaginary part gives the Hopf frequency:

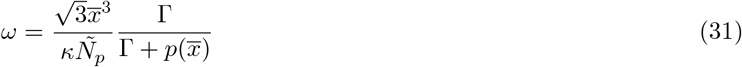

where the pre-factor is given by:

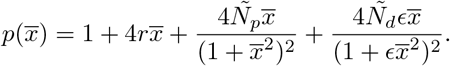

##### 2. Bifurcation analysis: ratio of frequency

We define *ω*_*r*_ to the ratio of Hopf frequencies with and without decoys for an given *N*_*p*_ (*ω* and *ω*_0_ respectively) while other parameters are equal. Using, Eq. 31, we get

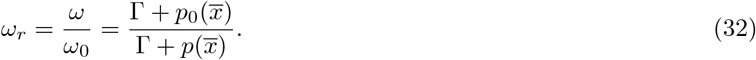

##### 3. Bifurcation analysis: condition for change in stability

The equilibrium point of the system will change its stability when the first pair of complex eigenvalues cross the imaginary axis. The point thus looses it stability and a limit cycle occurs via Hopf bifurcation.

We continue with Eq. 30 and equate the real part to obtain:

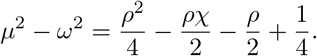

After substituting the expressions for *µ* and *ω*, and a few steps of algebra leads to the following expression:

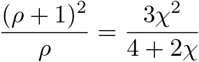

We again substitute the known expressions and identify Γ here as the critical Γ_*c*_, we find:

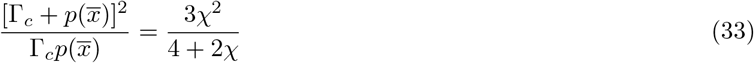

with 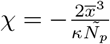.

Eq. 33 is quadratic in Γ and hence, generates two branches of the solution manifold dividing the parameter space in stable and unstable regions.

#### B. Parameters

We use the following parameters to perform analysis:

*C* = 10^−9^*M, c*_*p*_ = 10^7^*M* ^−1^, *c*_*d*_ = 10^7^*M* ^−1^, 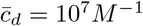, *σ* = 0.5.

|  |  |
| --- | --- |
| Time evolution of monomer concentration Fig. 4B | $\alpha = 9.60 \text{ min}^{-1}, \gamma_p = 0.08 \text{ min}^{-1}, \gamma_m = 0.10 \text{ min}^{-1}$ ($\Gamma = 0.80$ and $\kappa = 600.0$)[1-3] |
| Time evolution of monomer concentration Fig. 4C-D | $\alpha \approx 1.40 \text{ min}^{-1}, \gamma_p = 0.24 \text{ min}^{-1}, \gamma_m = 0.02 \text{ min}^{-1}$ ($\Gamma = 12.0$ and $\kappa = 145.0$)[1-3] |
| Frequency ratio and amplitude ratio variation Fig. 6 | $\alpha = 6.40 \text{ min}^{-1}, \gamma_p = 0.40 \text{ min}^{-1}, \gamma_m = 0.01 \text{ min}^{-1}$ ($\Gamma = 40.0$ and $\kappa = 800.0$)[1-3] |

---

**FIG 1.**
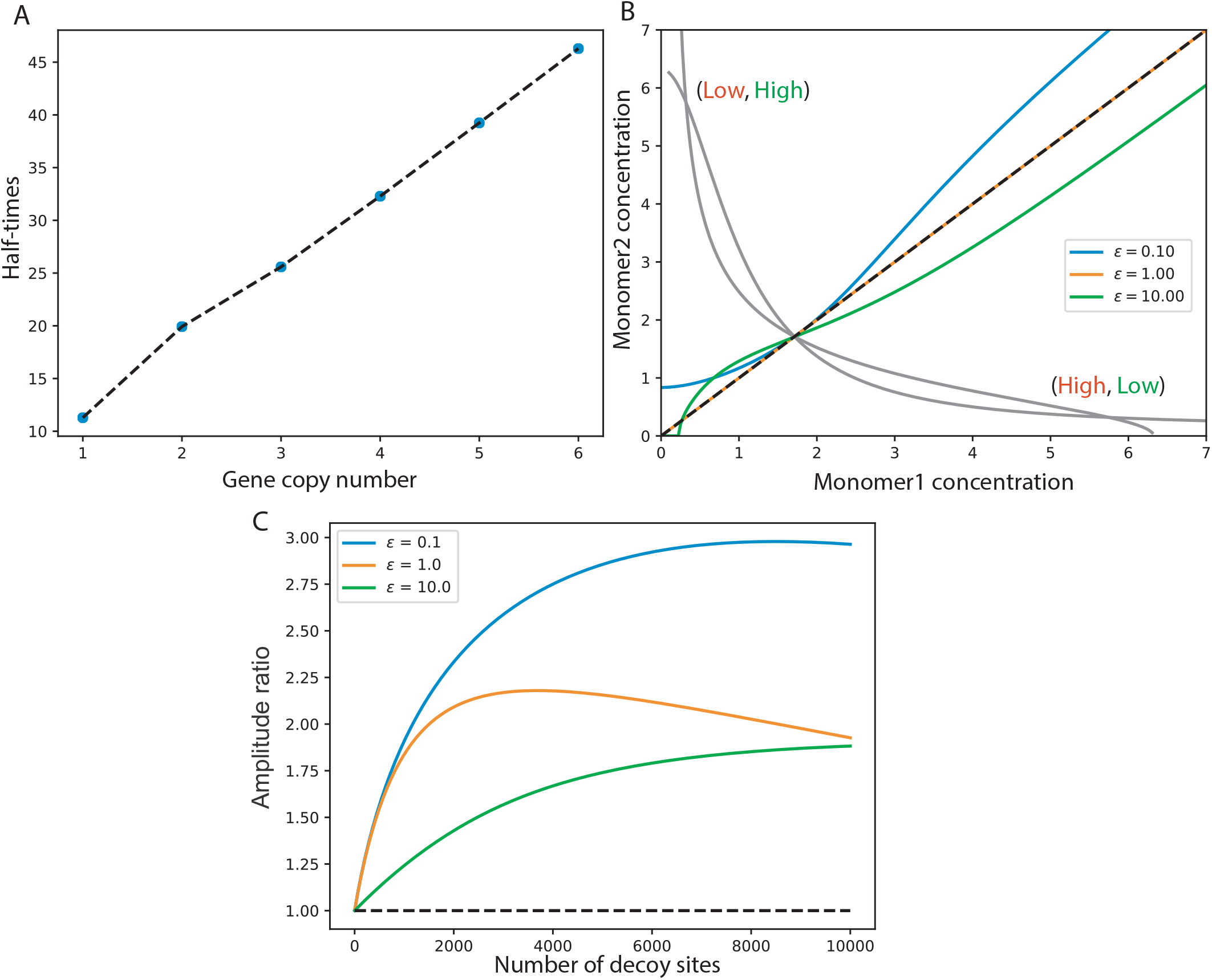
(A) The curves show linear increase in half-times to reach the steady-state with increase in gene copy number *N*_*p*_. For computational reliability, these half-times are computed by averaging over an ensemble of 100 random initial conditions chosen in the neighborhood of steady-state. (B) Separatices have been plotted for *N*_*p*_ = 1 for different protein binding affinities of decoys. The curve’s change is significant and therefore, basins of attraction of the two stables states are considerably altered. Amplitude ratio is highly sensitive to variation in protein binding affinities of decoys for *N*_*p*_ = 1. The effect appears to be suppressed for high binding affinities.

**FIG 2.**
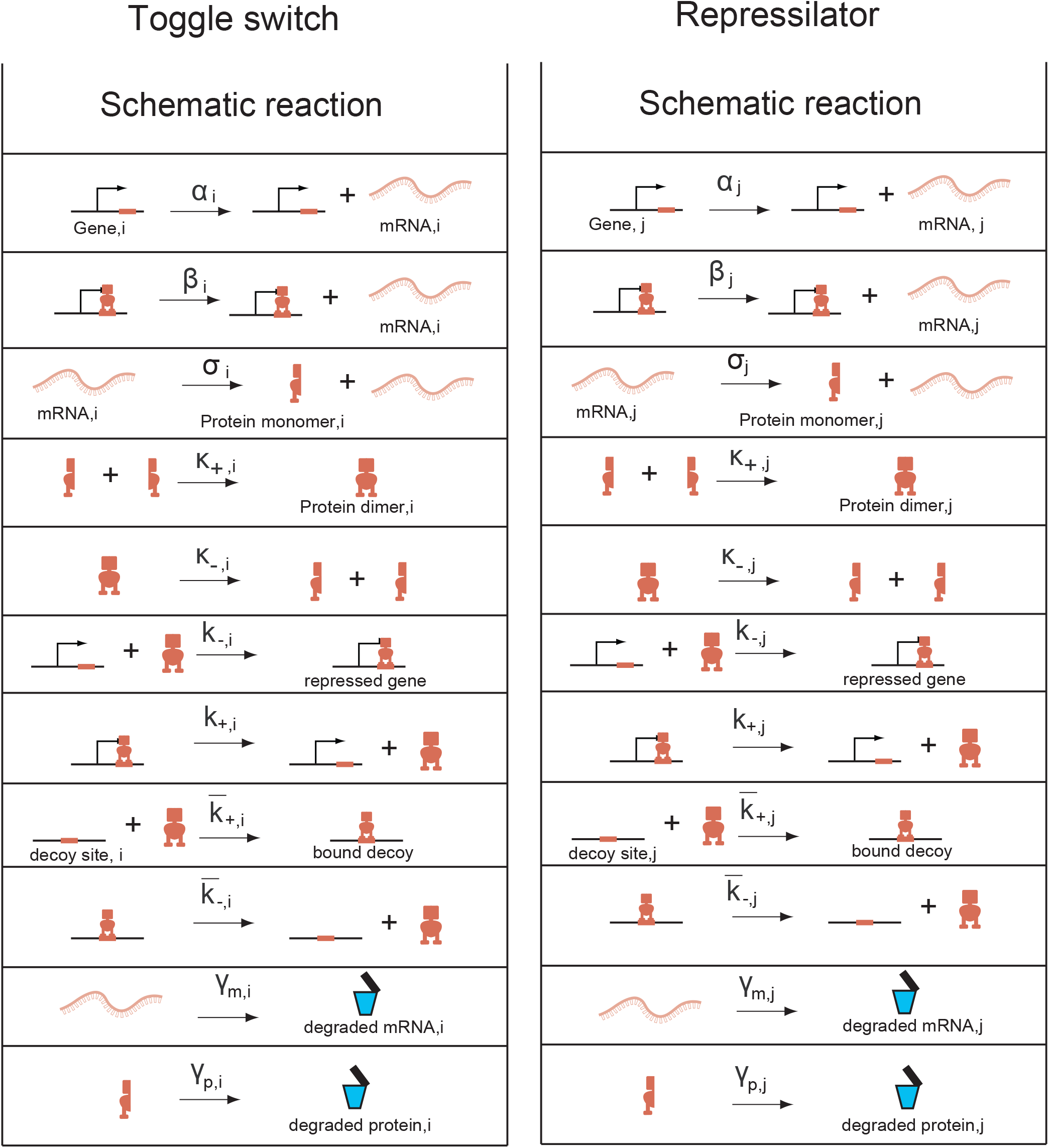
List of reactions used for toggle switch and repressilator. For toggle switch *i* takes values 1 and 2 corresponding to two genes of the motif. For repressilator *j* takes values 1, 2, and 3 corresponding to three genes of the motif.

**FIG 3.**
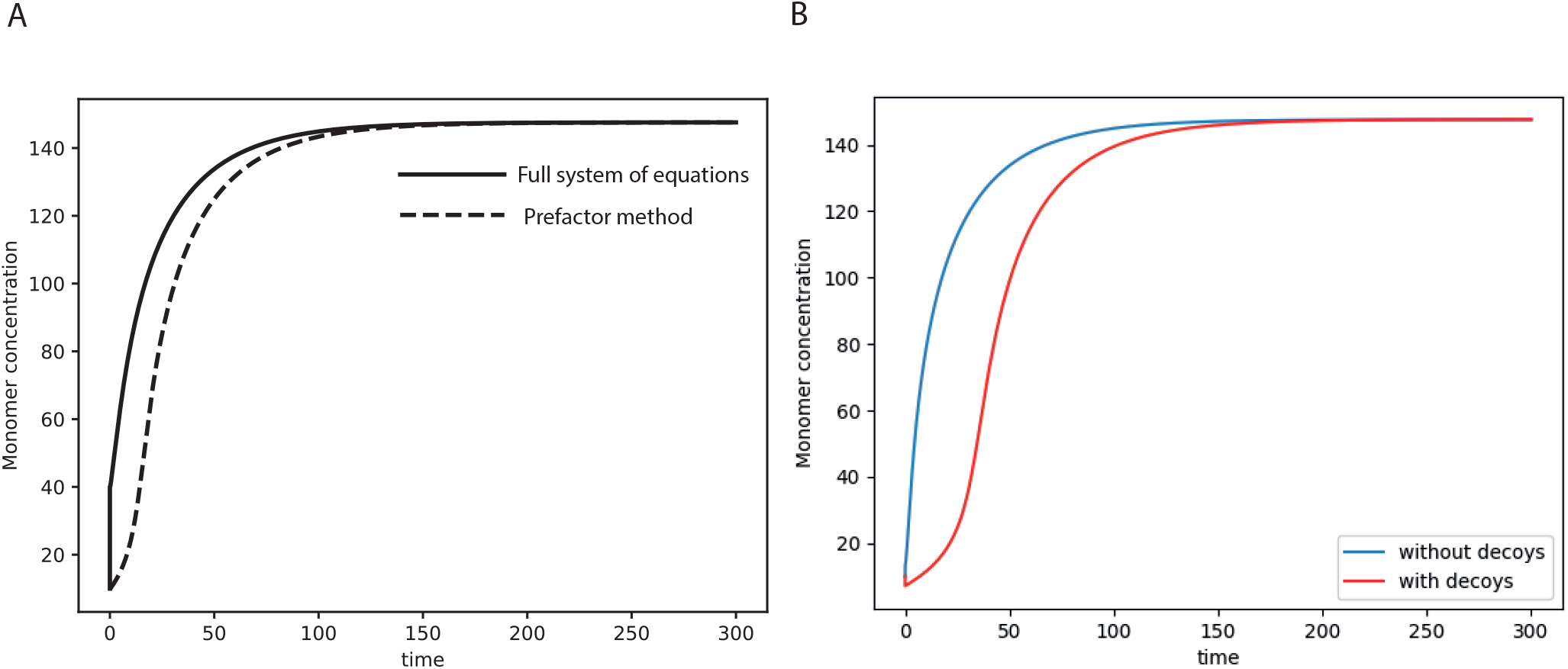
(A) Comparison of the pre-factor method (dashed curve) with the full un-reduced system (solid curves). (B) Transient dynamics due to increasing deocy sites sustains for longer times in the full system. The blue curve is for the case with decoy sites and the red curve shows the case with decoy sites. This behavior is again similar to what we have observed in the pre-factor method

**FIG 4.**
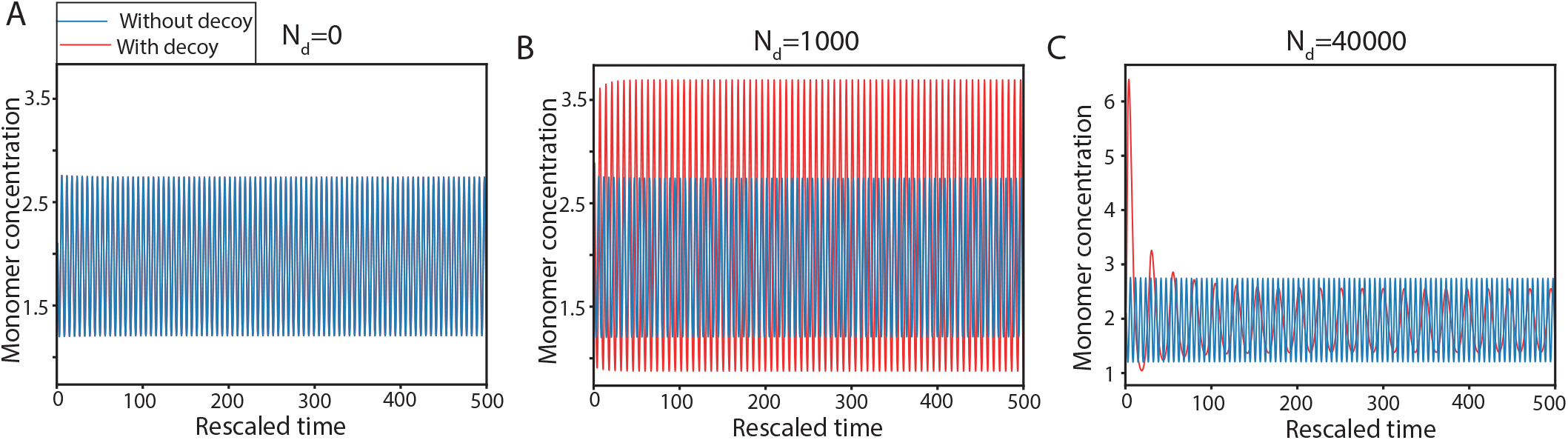
Examples to show variation in amplitude of Monomer concentration with time for different number of decoy sites at *N* = 1. The blue curve indicates the variation in the absence of decoy sites while the red curve shows that in the presence of decoy sites. (A) The case for the absence decoys i.e. *N*_*d*_ = 0.0. (B) For *N*_*d*_ = 1000, amplitude of oscillation increases. (C) Amplitude decreases at *N*_*d*_ = 40000.

